# Root anatomical gradients and cultivar differences underlie variation in root hydraulic properties in German winter wheat

**DOI:** 10.1101/2025.11.19.689226

**Authors:** Juan C. Baca Cabrera, Dylan H. Jones, Jan Vanderborght, Dominik Behrend, Hannah M. Schneider, Guillaume Lobet

## Abstract

Root hydraulic properties affect water uptake in wheat (*Triticum aestivum* L.) and are strongly influenced by root anatomy, yet how they vary along root axes and interact with cultivar differences remains underexplored. We investigated crown roots of six German winter wheat cultivars spanning one century of release, sampled from a field experiment. Roots were imaged at different positions along their axis using a high-throughput system (Rapid Anatomics Tool), and the resulting anatomical traits were coupled to GRANAR–MECHA to model radial (*K*_r_) and axial conductance (*k*_x_).

Longitudinal anatomical gradients were pronounced: tissue dimensions, metaxylem number, and apoplastic barriers decreased from the base onwards, resulting in *K*_r_ increasing and *k*_x_ decreasing with distance from the base. Cultivar differences were also apparent: modern cultivars had smaller tissues and fewer metaxylem vessels, reducing both axial and radial conductance and lowering whole-root water uptake capacity (∼20–30%).

By integrating field sampling with high-throughput image analysis and mechanistic modeling, this study establishes an integrated phenotyping approach that links root anatomy to water uptake and uncovers anatomical traits relevant to hydraulic function. The results show that longitudinal gradients and cultivar-associated anatomical differences contribute to variation in hydraulic properties and persist along fully mature root segments.

**Highlight:** High-throughput imaging–modeling shows that longitudinal gradients and cultivar-associated anatomical differences along crown roots shape radial and axial conductance, leading to reduced whole-root water uptake capacity in modern winter wheat

## Introduction

Wheat (*Triticum aestivum* L.) is a major global staple, providing roughly one-fifth of human calories and proteins and playing a central role in global food security (Erenstein *et al*., 2022). With climate change and declining freshwater availability projected to intensify the frequency and severity of droughts, thereby threatening agricultural production (Madadgar *et al*., 2017), improving the capacity of crops to access and use water efficiently has become increasingly urgent. Targeting root traits in breeding has been proposed as a promising strategy to develop more resilient and resource-efficient crops (Lynch, 2007). Root systems are essential for water and nutrient uptake and thus for sustaining crop growth and productivity (Torres-Ruiz *et al*., 2024), yet most breeding programs have historically focused on aboveground traits. This bias largely reflects the technical challenges of root phenotyping under field conditions (Atkinson *et al*., 2019) and the high plasticity of root traits (Schneider and Lynch, 2020).

In wheat, as in other cereal crops, root anatomy plays a pivotal role in water and nutrient transport and overall plant function (Steudle, 2000; Lynch *et al*., 2021). Traits such as cortex thickness, stele diameter, proportion of aerenchyma lacunae in the cortex, and the number and size of metaxylem vessels vary substantially between genotypes and along the root axis. (Yamauchi *et al*., 2019; Ouyang *et al*., 2020; Guhr *et al*., 2025), but these longitudinal patterns and their interaction with cultivar differences have not been quantified in a comprehensive way. Monocot roots grow from the root apex, dynamically producing new transport tissue as the root elongates. However, they lack the capacity for radial secondary growth, so that following differentiation, this initially formed tissue is what persists throughout the life of the root (Wu *et al*., 2011; Clément *et al*., 2022; Petrova *et al*., 2023). Both developmental gradients and local environmental conditions shape variation in cortex expansion, vascular differentiation, and apoplastic barrier formation (Jones *et al*., 2025) —key determinants of hydraulic function.

Anatomical traits influence both the radial (movement of water into the root) and axial (movement of water along the root axis) components of root water flow (Chimungu *et al*., 2014; Lynch *et al*., 2014; Schneider *et al*., 2017; Cuneo *et al*., 2021; Yamauchi *et al*., 2021). In particular, the formation of apoplastic barriers, such as suberization and Casparian strips in the endodermis, strongly increases radial resistance (Geldner, 2013; Song *et al*., 2023). As these traits change along the root axis, axial conductance (*k*_x_) and radial conductivity (*k*_r_) are expected to vary, as has been shown in wheat (Bramley *et al*., 2009). Yet, experimental studies quantifying these properties remain scarce, largely due to the technical complexity of segment-scale hydraulic measurements (Boursiac *et al*., 2022*b*). Recent advances in high-throughput, low-cost anatomical phenotyping (Jones *et al*., 2025, Preprint), combined with explicit modelling of water transport at the cell scale (Couvreur *et al*., 2018), provide new opportunities for more systematic and efficient analysis of root hydraulic properties, beyond what is feasible with purely experimental approaches.

In a previous study on historical German winter wheat cultivars released between 1895 and 2002, we observed a significant decline in whole root system conductance (*K*_rs_) and root axis number with cultivar release year (Baca Cabrera *et al*., 2025). Given that *K*_rs_ reflects the maximum root water uptake capacity under sufficiently wet soil conditions, this trend suggested that breeding may have inadvertently favoured more conservative root water-uptake strategies, i.e., a water-saving behaviour aimed at maintaining plant water status when water availability becomes limiting (Richards and Passioura, 1989; Blessing *et al*., 2018). Root architectural phenotyping revealed that the decline in *K*_rs_ was associated with a reduction in root axis number. However, whether anatomical changes also contributed remains unknown, as anatomy was not assessed in that study. Furthermore, it remains unclear whether the effect of breeding on *K*_rs_ was driven primarily by changes in *k*_r_, *k*_x_, or both. Thus, understanding how root hydraulic properties have evolved with breeding is critical for explaining variation in water uptake strategies in modern cultivars.

Building on this, here we investigated longitudinal gradients and cultivar differences in wheat root anatomical and hydraulic traits, using the same six historical German winter wheat cultivars previously analyzed in Baca Cabrera *et al*. (2025). Specifically, we focused on crown roots, the dominant axes for water and nutrient uptake in cereals (Steffens and Rasmussen, 2016). Our study addressed three main questions: (i) do root anatomical traits (cortex, stele, and metaxylem dimensions; proportion of aerenchyma lacunae in the cortex; and apoplastic barrier development) vary along the root axis?; (ii) are these traits and their interaction with longitudinal gradients related to the cultivar release year; and (iii) what is the effect of anatomical variation on root hydraulic properties (*k*_r_ and *k*_x_) and its implications for root water uptake capacity? To answer these questions, we developed an integrated phenotyping approach that combined detailed anatomical measurements using the Rapid Anatomics Tool (RAT) (Jones *et al*., 2025, Preprint) with root hydraulic modelling through the GRANAR–MECHA framework (Heymans *et al*., 2019), enabling us to link root structure and function.

## Materials and Methods

### Plant material and root sampling

Plant material was obtained from a field experiment that has been previously described in detail in Baca Cabrera *et al*. (2025) and Behrend *et al*. (2025, Preprint). In brief, six German winter wheat cultivars (*Triticum aestivum* L.) spanning more than 100 years of breeding history were grown under rainfed conditions at the Campus Klein-Altendorf research station near Bonn, Germany (50°37’ N, 6°59’ E) during two consecutive growing seasons (2022–2023 and 2023–2024). The soil at the site is a Haplic Luvisol developed on loess, known to be very homogeneous. Field management followed standard agronomic practices with conventional fertilization and weed control, and there was no sign of water stress during growth, with precipitation above the long-term average in both growing seasons (Supplementary Fig. S1).

The cultivars were selected based on historical relevance, availability, and consistency with previous studies, and their release years were spaced at approximately 20-year intervals: S. Dickkopf (1895), SG v. Stocken (1920), Heines II (1940), Jubilar (1961), Okapi (1978), and Tommi (2002). All cultivars were grown in a randomized block design with four field replicates.

Sampling took place at the end of the tillering stage (BBCH < 30) during both growing seasons (spring 2023 and spring 2024), with all cultivars harvested simultaneously in single-day campaigns. Root samples were collected using a slightly modified “shovelomics” method for wheat (York *et al*., 2018, Preprint). From each plot, a representative topsoil portion (20–30 cm deep, ∼30 cm diameter) was excavated, typically yielding 5–10 plants per sample. Samples were stored at 5 °C until root washing. In the laboratory, roots were soaked in water and carefully washed, and root crowns were separated from the shoots close to the base (leaving ∼3 cm of tiller tissue). Samples of 4 plants per plot were preserved in a water (37.5%)–ethanol (37.5%)–glycol (25%) solution for further analyses. From these samples, the longest crown root from each plant (∼20–25 cm long) was excised directly at the tiller junction (root base) and stored in vials for subsequent anatomical imaging, resulting in a total of 32 crown roots per cultivar (*n* = 32).

### Root subsampling and anatomical imaging

From each selected crown root, 2–3 cm segments were excised from three positions: basal (the 2.5 cm closest to the tiller junction), mid-segment (approximately 10 cm from the base), and distal (approximately 3 cm from the most distal portion of the root, i.e., ∼20 cm from the base).

Subsamples were stored in 2 mL Eppendorf microcentrifuge tubes filled with 70% ethanol (v/v) at 4 °C for a minimum of two weeks. It is worth noting that due to the sampling protocol, the root tip (meristematic and elongation zones) was not included.

Cross-section imaging of crown roots was performed using the Rapid Anatomics Tool (RAT) method as described in Jones *et al*. (2025, Preprint), which has been tested on a range of diverse plant samples, and delivers reliable, high-quality anatomical images (Supplementary Fig. S2). This approach combines blockface-like imaging, where a fresh surface of the root is exposed for imaging, with stain-free near-ultraviolet (nUV) induced autofluorescence to highlight cell walls. In brief, preserved root segments were mounted in a 3D-printed holder to ensure consistent orientation and positioning, then cut to reveal a clean cross-section. Imaging was performed using a USB microscope (Dino-lite Edge AM8517MT-FUW) with nUV illumination at 220x magnification. Minor adjustments in fluorescence contrast were applied when necessary to improve tissue visualization. Images were analyzed in Fiji (ImageJ) (Schindelin *et al*., 2012) to quantify the metaxylem (number, diameter and area), stele (diameter and area), cortex (cortical cell thickness and file layer number and total cortex area), aerenchyma (proportion of the cortex occupied by aerenchyma lacunae), and whole-root cross-section (diameter and area). Stele and cortex diameters were measured at their narrowest points using the line tool, areas were measured manually using the polygon tool. For metaxylem, images were converted to 8-bit, and a 2-pixel radius Gaussian blur was applied (pixel size: 1.365 μm). The magic wand tool was set to a threshold of 20 (adjusted ad-hoc) to select the area of each metaxylem lumen individually.

To complement the anatomical measurements, several tissue ratios were calculated to capture allometric relationships. Specifically, we computed the cortex-to-stele ratio (CSR, cortex area ÷ stele area), stele-to-root ratio (SRR, stele area ÷ total root area), xylem-to-stele ratio (XSR, total metaxylem area ÷ stele area), xylem-to-root ratio (XRR, total metaxylem area ÷ total root area), and a combined index (XCS), defined as (total metaxylem area × cortex area) ÷ (stele area^2^). These ratios have previously been used to assess anatomical investment patterns in grasses (Yamauchi *et al*., 2021) and facilitate the comparison of tissue allocation across cultivars and root positions (Jones *et al*., 2025, Preprint).

Additionally, to qualify the presence of apoplastic barriers (specifically hypodermal modifications) in the root anatomical image set, images were individually evaluated and scored. The range of images were initially appraised for presence or absence of hypodermal modifications as described in Cantó-Pastor *et al*. (2025) and Jones *et al*. (2025). Lignin distribution could be assessed from images captured using nUV autofluorescence (Schneider *et al*., 2021; Hazman and Kabil, 2022; Cunha Neto *et al*., 2023). Three categories of hypodermal modification were observed, presence of multiseriate sclerenchyma, presence of a differentiated exodermis (either polar, Y-shaped, polar, or anticlinal), and a combination thereof. Two other categories were assigned, the absence of hypodermal differentiation (relative to cell walls of the cortical mesodermis), and cortical senescence (Supplementary Table S1). Where there was variation within an image or disruption on the cortex caused by lateral root emergence or damage, values were assigned based on the largest contiguous non-damaged region. In all images a differentiated endodermis with Casparian strip was visible, typically stage III (U-shaped). All anatomical measurements were performed at the Leibniz Institute of Plant Genetics and Crop Plant Research (IPK). Raw data processing was performed in R to obtain anatomical traits for each image, at each root position.

### Modelling of segment-scale root hydraulic properties

Segment-scale axial conductance (*k*_x_, m⁴ MPa⁻¹ s⁻¹) and radial conductivity (*k*_r_, m MPa⁻¹ s⁻¹) and conductance (*K*_r_, m^3^ MPa^-1^ s^-1^) were estimated using the GRANAR–MECHA computational framework, which predicts emergent root hydraulic properties from anatomical cross-sections. For each crown root at each sampling position, root anatomies were generated with the GRANAR (v1.1) model (Heymans *et al*., 2019), based on measured anatomical traits (root and stele diameter, cortical cell file layer number and thickness, and number and diameter of metaxylem vessels). GRANAR builds virtual cross-sections by placing cell layers around the root center according to the input traits, introducing a small random variation in cell positions to create unique anatomies while preserving overall tissue features. The reconstructed anatomies (Supplementary Fig. S3) were then processed with MECHA (Couvreur *et al*., 2018) to model radial water flow and *k*_r_, using default subcellular hydraulic parameters (Supplementary Table S2).

MECHA can simulate alternative apoplastic barrier scenarios, which strongly affect radial water flow and *k*_r_ (Couvreur *et al*., 2018). The model includes nine scenarios defined by the presence or absence of a Casparian strip (CS) and/or full suberization (Sub) in the endodermis and exodermis: (0) no barriers, (1) endodermis with CS, (2) partially suberized endodermis with passage cells, (3) fully suberized endodermis, (4) fully suberized endodermis + exodermis with CS, (5) endodermis with CS + exodermis with CS, (6) endodermis with CS + fully suberized exodermis, (7) exodermis with CS only, and (8) fully suberized endodermis + fully suberized exodermis. In this study, the barrier scenario assigned to each cross-section was matched to the observed presence of apoplastic barriers, according to our scoring system.

Radial conductance (*K*_r_) was further calculated as

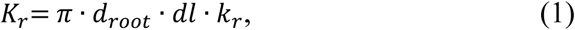

where *K*r is the radial conductance of a root segment of infinitesimal length *dl* (m), *d*_root_ is the root diameter (m), and *k*_r_ is the radial conductivity. *k*_x_ was calculated within MECHA using the Hagen–Poiseuille law, based on the number and diameter of mature metaxylem vessels, while protoxylem elements were not included in the calculation.

In total, 573 root cross-sections were processed. Of these, >99% could be successfully reconstructed with GRANAR, and >92% of the reconstructions could be modeled with MECHA, underscoring the robustness of the workflow and the representativeness of the obtained root hydraulic properties across cultivars and root positions. This workflow provides an integrated pipeline for quantifying root anatomical and hydraulic traits from field grown plants (Fig. 1).

**Figure 1:**
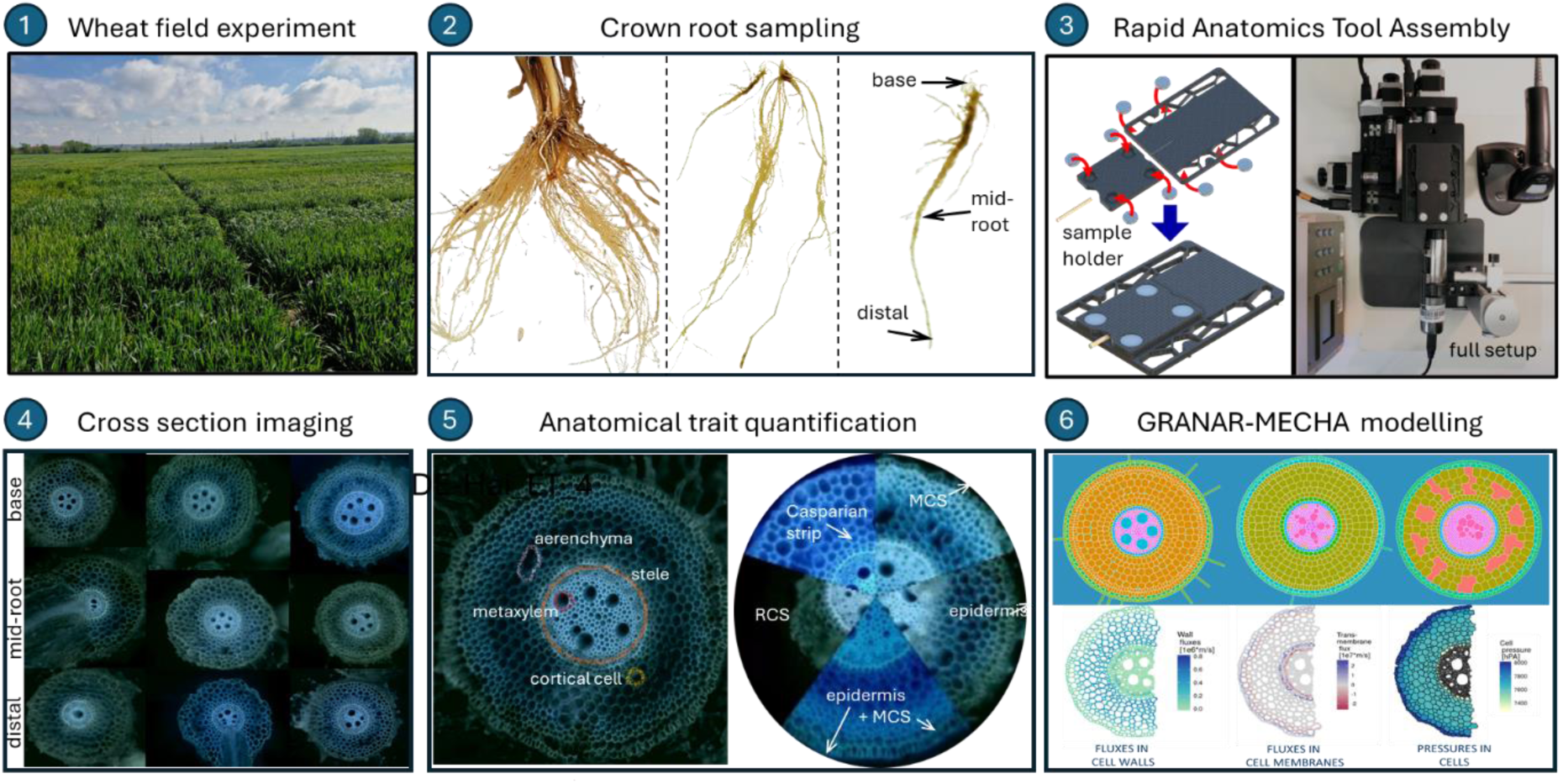
Integrated phenotyping approach for the quantification of root hydraulic properties. Crown roots were sampled from a field experiment (Behrend *et al*., 2025, Preprint) and 2–3 cm root segments at the base, mid-root and distal positions were subsampled for cross section imaging using the Rapid Anatomics Tool (RAT) (Jones *et al*., 2025, Preprint). Traits obtained from the images were used as input for the quantification of root hydraulic properties using the GRANAR-MECHA modelling framework (Heymans *et al*., 2019).

### Statistical analysis

All statistical analyses were conducted in R v.4.4.1 (R Core Team, 2024). Effects of cultivar year of release and root position, as well as their interaction, were assessed using two-way ANOVA, with model choice depending on the trait type. Continuous variables (i.e., tissue diameters, areas, and axial conductance), anatomical ratios (XSR, CSR, XRR, SRR, XCS), and count variables (number of cortical cell file layers and metaxylem vessels) were analyzed with ordinary least-squares linear models. Continuous variables and ratios were log-transformed before model fitting, while count data were analyzed untransformed. Radial conductivity was analyzed using a rank-based linear model (Kloke and McKean, 2012) due to discontinuities caused by large differences in radial water flow associated with the presence of apoplastic barriers. Aerenchyma, being strongly zero-inflated and expressed as a ratio, was analyzed using a generalized linear model with a quasibinomial link. Model validation was performed by visual inspection of residuals (Q–Q plots). For plotting purposes, all transformed traits were back-transformed to their original scale. Additionally, apoplastic barrier presence was analyzed using multinomial regression (nnet package; Venables and Ripley, 2002).

To explore overall patterns of anatomical variation, a principal component analysis (PCA) was performed on directly measured anatomical traits (excluding derived or modelled variables, and apoplastic barrier scoring) with the package FactoMineR (Lê *et al*., 2008), followed by K-means clustering of trait loadings. The ggplot2 package (Wickham, 2016) was used for data visualization.

## Results

### Root anatomical traits across root positions and cultivar release year

To quantify anatomical variation along crown roots, cross-sections were imaged at basal, mid, and distal positions (Supplementary Fig. S2). Across cultivars, these analyses revealed clear longitudinal gradients in tissue dimensions and metaxylem structure (Fig. 2A–E, Table 1). All measured traits differed significantly among root positions (*p* < 0.001; Fig. 2A–E, Table 1), with all decreasing from the base toward the tip, except for cortical cell file layer diameter, which showed the opposite trend. Across all cultivars, the largest variation along the root was observed for aerenchyma, with a 59.5% decrease in the proportion of the cortex occupied by aerenchyma lacunae, from the basal to the distal sampling point. The smallest differences were found in cortical tissues, with cortical cell file layer diameter increasing by 7.4%, while cortical cell file layer number and total cortex thickness decreased by 14.9% and 12.5%, respectively. Metaxylem area and stele area decreased by similar magnitudes from the base toward the tip (33.8% and 39.4%, respectively), leading to a corresponding 24.5% decline in total root area along the root axis.

**Figure 2:**
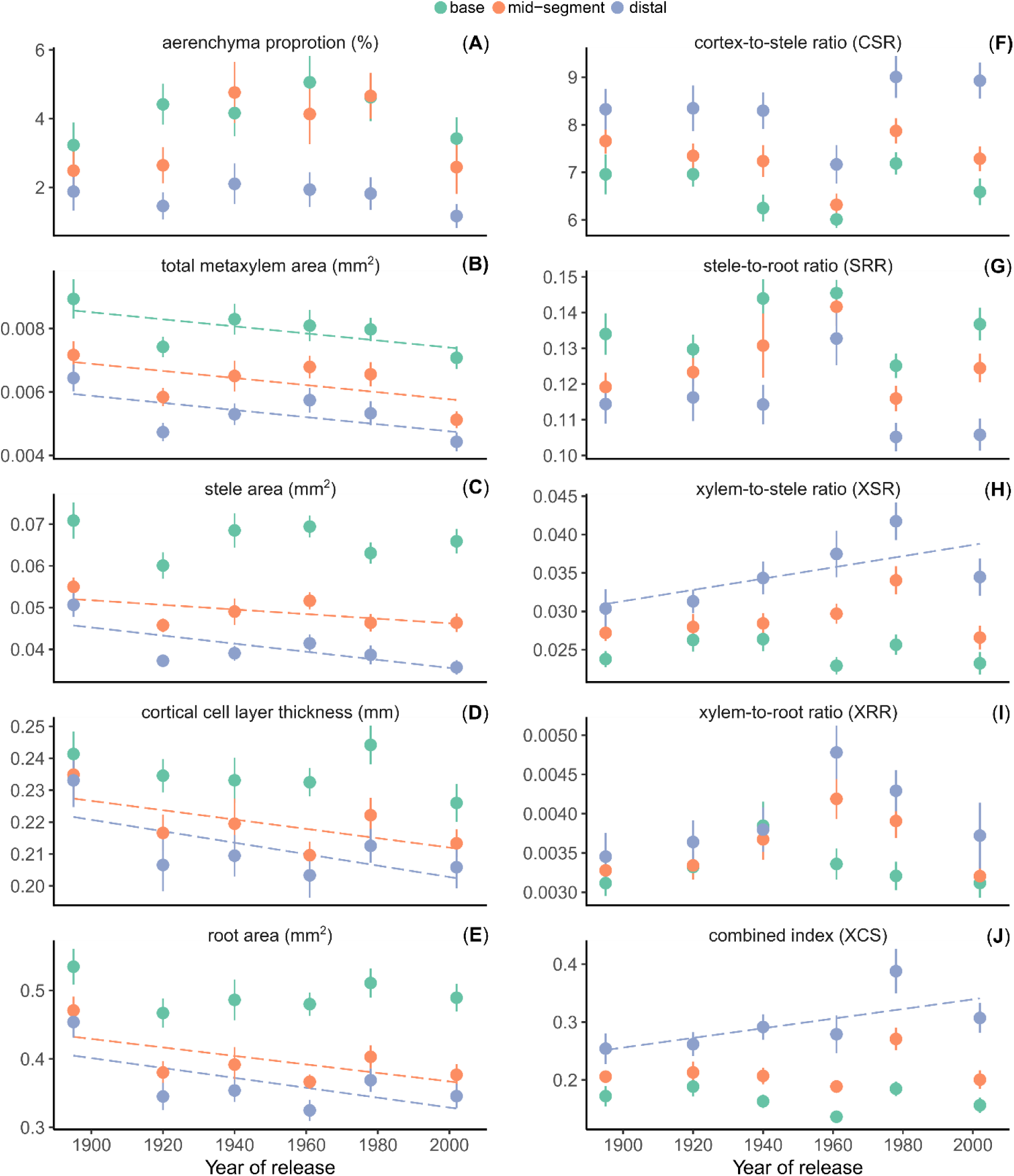
Root anatomical traits at three positions along crown roots (base, mid-segment and distal) for 6 different cultivars of winter wheat (*T. aestivum* L.). Panels (**A**) to (**E**) show traits obtained from cross section images, and panels (**F**) to (**J**) derived allometric ratios (cortex-to-stele area, stele-to-root area ratio, metaxylem-to-stele area ratio, metaxylem-to-root area ratio, and composite index XCS). Data points and error bars represent the mean ± SE (*n* = 32). Dashed lines represent the linear relationship between cultivar release year and the analyzed traits at each root position (only shown if significant, *p* < 0.05).

**Table 1:** Root traits of winter wheat (*T. aestivum* L.), as influenced by cultivar release year, root position and their interaction. Effect significance (*p*-value) and trait range (average values) across all combinations of cultivars (six German cultivars) and root positions (basal, mid-root and distal) (*n* = 32 per group).

| trait | range | *p*-value |  |  |
| --- | --- | --- | --- | --- |
|  |  | release year | root position | release year × position |
| root area (mm<sup>2</sup>) | 0.32 – 0.53 | <0.001 | <0.001 | 0.11 |
| metaxylem area (mm<sup>2</sup>) | 0.0011 – 0.0017 | 0.21 | <0.001 | 0.83 |
| metaxylem number | 3.6 – 5.5 | <0.01 | <0.001 | 0.13 |
| total metaxylem area (mm<sup>2</sup>) | 0.0044 – 0.0089 | <0.001 | <0.001 | 0.57 |
| cortical cell file layer diameter (mm) | 0.026 – 0.031 | <0.01 | <0.001 | 0.99 |
| cortical cell file layer number | 7.2 – 9.2 | 0.57 | <0.001 | 0.36 |
| total cortex thickness (mm) | 0.20 – 0.24 | <0.01 | <0.001 | 0.66 |
| stele area (mm<sup>2</sup>) | 0.036 – 0.071 | <0.001 | <0.001 | <0.05 |
| aerenchyma proportion | 0.012 – 0.051 | 0.54 | <0.001 | 0.52 |
| $k_x$ (m<sup>4</sup> MPa<sup>-1</sup> s<sup>-1</sup> × 10<sup>-9</sup>) | 0.56 – 1.6 | <0.01 | <0.001 | 0.68 |
| $k_r$ (m MPa<sup>-1</sup> s<sup>-1</sup> × 10<sup>-7</sup>) | 0.67 – 1.1 | 0.47 | <0.001 | 0.78 |
| $K_r$ (m<sup>3</sup> MPa<sup>-1</sup> s<sup>-1</sup> × 10<sup>-7</sup>) | 1.6 – 2.5 | <0.05 | <0.001 | 0.91 |
| XSR | 0.023 – 0.042 | 0.06 | <0.001 | <0.05 |
| CSR | 6.0 – 9.0 | 0.81 | <0.001 | 0.38 |
| XRR | 0.0031 – 0.0048 | 0.13 | <0.001 | 0.17 |
| SRR | 0.11 – 0.15 | 0.83 | <0.001 | 0.36 |
| XCS | 0.14 – 0.39 | 0.13 | <0.001 | <0.05 |

Cultivar year of release (YOR) had a significant effect (*p* < 0.01) on total metaxylem area, stele area, total cortex thickness, and root area, but not on the proportion of the cortex occupied by aerenchyma lacunae (Fig. 2A–E, Table 1). In all significant cases, the effect indicated smaller tissue areas with increasing YOR. Averaged across root sampling positions, stele area decreased from 0.059 mm² in the oldest cultivar to 0.049 mm² in the most modern cultivar, and a similar decline was observed for root area (from 0.49 mm² to 0.40 mm²) (Table S3). The reduction in metaxylem area with YOR was associated with a significant decrease in the number of metaxylem vessels (from ca. 5 to 4 between the oldest and most modern cultivars), while the area of individual metaxylem vessels did not vary significantly with YOR (*p* = 0.21). By contrast, the decrease in cortex thickness with YOR was associated with a significant reduction in cortical cell layer diameter (from 0.030 mm to 0.027 mm), but not in the number of cortical cell file layers (average of 8 across cultivars) (Table S3). In most cases, no significant interaction was observed between YOR and root position. The only exception was stele area (*p* < 0.05, Table 1), which did not vary among cultivars at the root base but was significantly smaller in mid-segment and distal positions of modern cultivars compared to older ones (Fig. 2c, Table S3).

Further, anatomical ratios were calculated to evaluate allometric relationships between tissues, including the metaxylem-to-stele area ratio (XSR), cortex-to-stele area ratio (CSR), metaxylem-to-root area ratio (XRR), stele-to-root area ratio (SRR), and the composite index XCS (Materials & Methods). Similarly to the individual traits, all ratios differed significantly among root positions (*p* < 0.001). However, the direction of change was the opposite: except for SRR, all ratios increased from the base towards the tip (Fig. 2F–J, Table 1). YOR did not have a significant main effect on the ratios, but significant interactions between root position and YOR were detected for XSR and XCS (*p* < 0.05). In both cases, the interaction reflected a shift at the distal position: while no differences among cultivars were observed at the base and mid-segment, modern cultivars exhibited significantly higher XSR and XCS values than older ones toward the tip, driven by a higher relative area of the stele occupied by metaxylem (Fig. 2).

### Principal component analysis of anatomical traits

A PCA was performed to further explore the variation in anatomical traits across root positions and cultivars (excluding allometric ratios, see Materials & Methods). The PCA captured the major structure of trait variation, with the first two axes explaining 69.3% of the total variance (50.3% and 19.0% for PC1 and PC2, respectively; Fig. 3A-B). Root position was the dominant source of variation: base, mid-segment, and distal samples formed three clearly separated groups along PC1 (Fig. 3A). Cultivar differences were less pronounced, although the oldest cultivar (S. Dickkopf) consistently clustered apart from the others towards higher values of both PC1 and PC2, while the remaining cultivars largely overlapped in ordination space (Fig. 3B).

**Figure 3:**
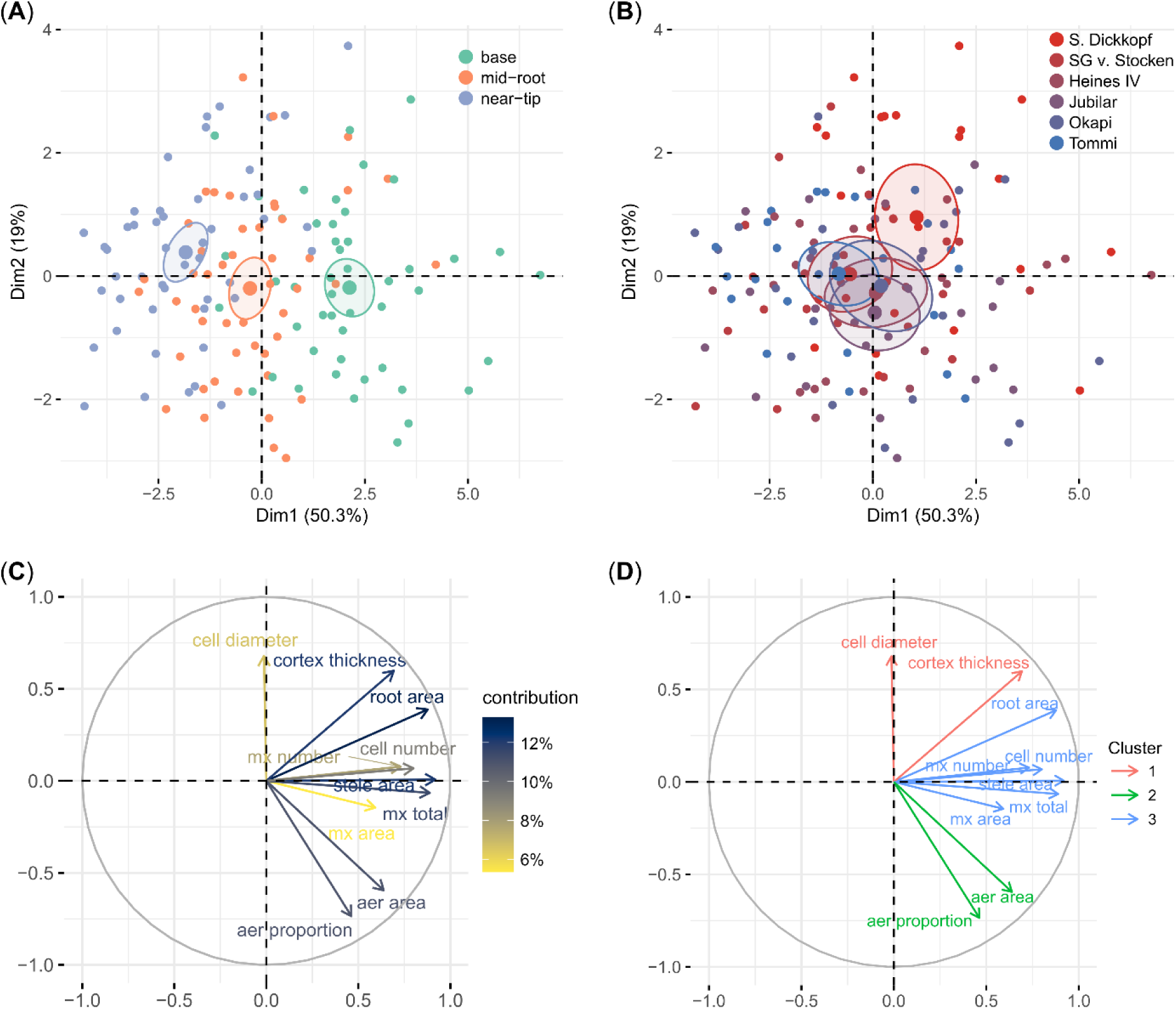
Principal component analysis (PCA) of root anatomical traits at three positions along crown roots (base, mid-segment and distal) for 6 different cultivars of winter wheat (*T. aestivum* L.). PCA scores, showing separation by root position (**A**) and by cultivars (**B**). PCA loadings indicating the contribution of each anatomical trait to the first two principal components (**C**). Clustering of traits into three groups using k-means, highlighting shared patterns of variation (**D**). Large symbols in (**A**) and (**B**) represent the grouping centroids, and ellipses indicate the 95% confidence intervals

Among all variables, tissue dimensions (stele, total metaxylem and root area and total cortex thickness) contributed most strongly to the ordination, whereas individual metaxylem vessel area and individual cortical cell file layer diameter showed the weakest contributions (Fig. 2C). Trait loading also indicated that PC1 was mainly associated with the tissue areas, while PC2 was influenced by cortical features and aerenchyma presence. Additionally, K-means clustering of trait contributions confirmed three coherent trait groups: (i) cortex-related traits (cell file layer diameter, cortex thickness), (ii) aerenchyma, and (iii) all remaining size-related traits (Fig. 2D). Size-related traits largely drove the separation among root positions, while cultivar differences were less pronounced and reflected both cortical traits and tissue size.

### Apoplastic barrier development across root positions and cultivars

Apoplastic barrier development was qualitatively assessed from the same anatomical images by scoring hypodermal modifications, including the presence of multiseriate cortical sclerenchyma (MCS), exodermis, exodermis with MCS, or cortical senescence (Scores 1–5; Materials & Methods; Table S1). In all sampled segments, even at the distal sampling position (∼20 cm from the base), the endodermis was already differentiated with a Casparian strip (and in some cases suberization) and thus does not contribute to variation in these scores. Therefore, variation in apoplastic barrier development in this dataset was primarily driven by hypodermal tissues.

The distribution of hypodermal apoplastic barrier states strongly depended on the root position (Fig. 4). At the distal position, the vast majority of samples (>75% across all cultivars) showed no hypodermal modification (Score 1). In mid-segment samples, this state remained dominant (>50%), but exodermis formation (Score 3) also became frequent (15–45%, depending on cultivar). At the base, the distribution was more balanced among categories, with exodermis plus MCS (Score 4, >40% of observations) being most common in the three most modern cultivars, endodermis only (Score 1) prevailing in the two oldest cultivars (>30%), and exodermis (Score 3, 45%) being most frequent in the remaining cultivar. Cortical senescence (Score 5) was rare and never observed at the root base.

**Figure 4:**
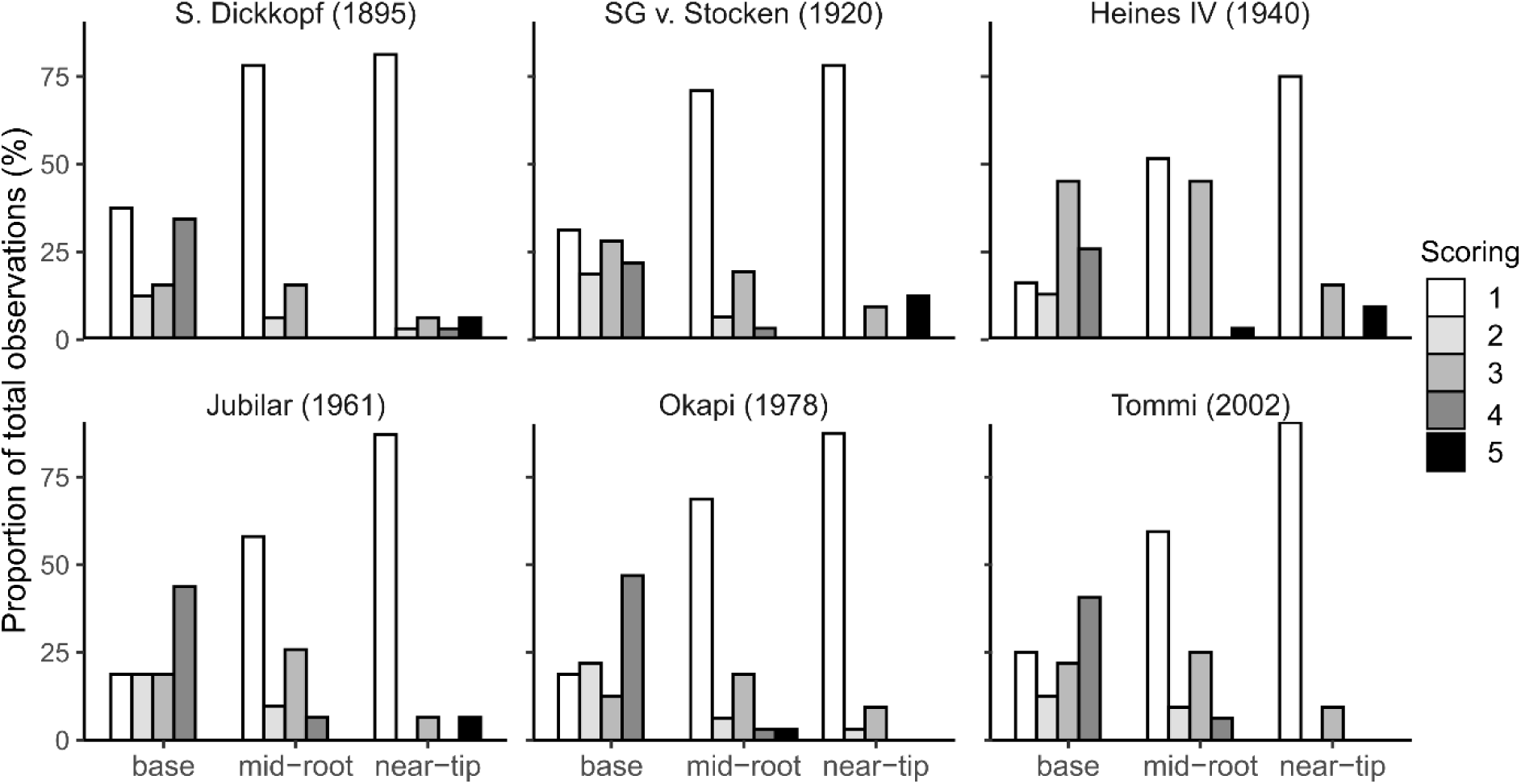
Distribution of apoplastic barrier development at three positions along crown roots (base, mid-segment and distal) for 6 different cultivars of winter wheat (*T. aestivum* L.). Apoplastic barrier development (defined as hypodermal modifications) was scored from anatomical cross-sections as: (1) no modifications, (2) multiseriate cortical sclerenchyma (MCS), (3) exodermis, (4) exodermis and MCS combined, or (5) cortical senescence. Bars show the relative frequency of scores within each cultivar × root position group (*n* = 32 per group).

Multinomial regression confirmed that apoplastic barrier status varied significantly among root positions (likelihood ratio vs. null model, *p* < 0.001), whereas cultivar release year and its interaction with root position were not significant (*p* > 0.05). This indicates that, although some differences among cultivars may appear, no consistent trend in barrier development could be detected with increasing YOR.

### Root hydraulic properties development along the root axis

Axial conductance (*k*_x_) and radial conductivity (*k*_r_) were estimated for each cultivar and root position using the GRANAR–MECHA modelling pipeline (Materials & Methods). Both traits showed highly significant differences among root positions (*p* < 0.001; Fig. 5A–B, Table 1). Meanwhile, cultivar release year had a significant effect on *k*_x_ (*p* < 0.05), but not on *k*_r_, and no interaction between root position and release year was observed.

**Figure 5:**
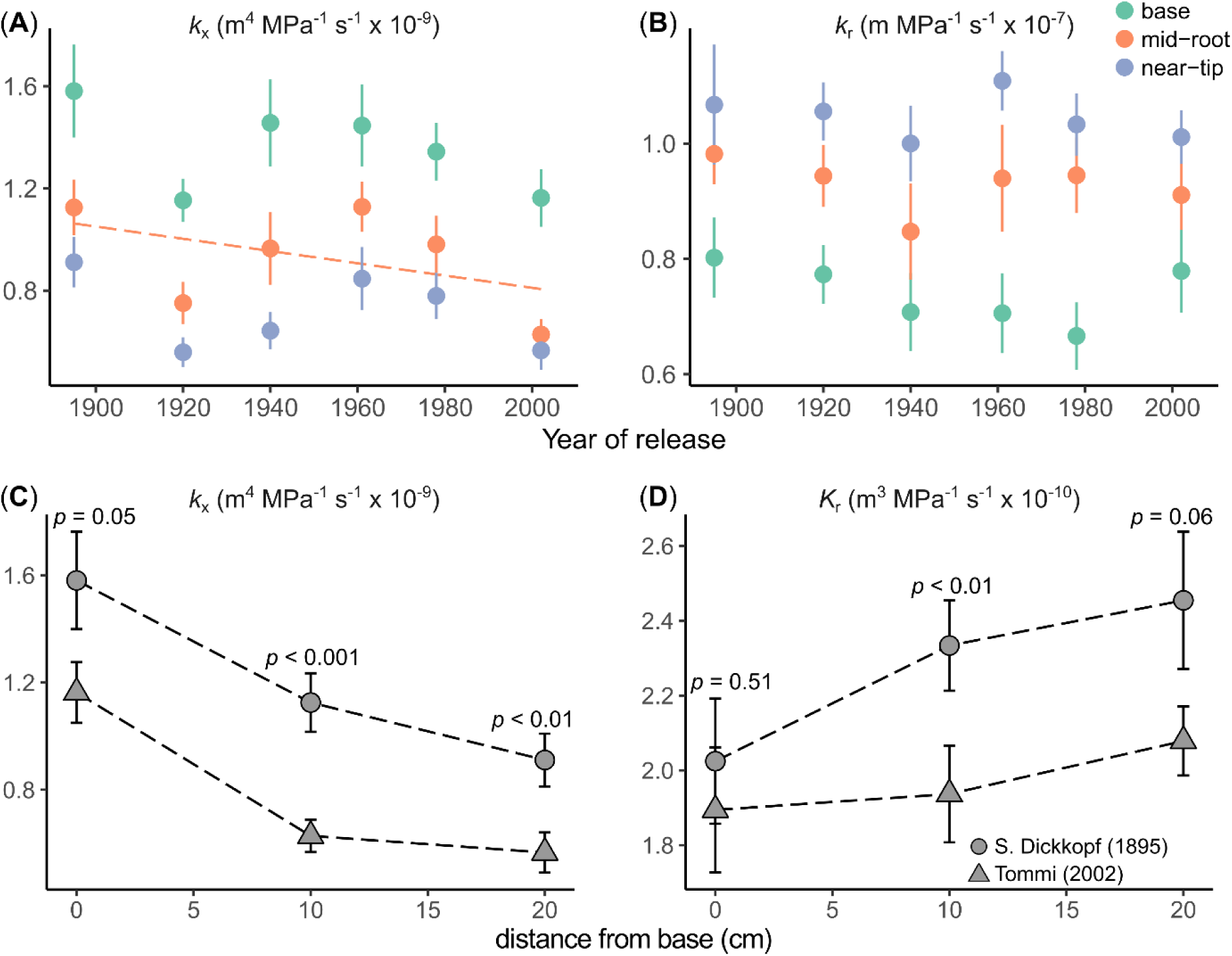
Root hydraulic properties of winter wheat cultivars (*T. aestivum* L.). (**A**–**B**) Axial conductance (*k*_x_) and radial conductivity (*k*_r_) at three positions along crown roots (base, mid-segment and distal) for 6 different cultivars. (**C**–**D**) Spatial dynamics of *k*_x_ and radial conductance (*K*_r_, i.e. *k*_r_ scaled by surface area) along the root axis, illustrated by comparing the oldest and most modern cultivars. Data points and error bars represent the mean ± SE (n = 32). Dashed lines in (**A**) and (**B**) represent the linear relationship between cultivar release year and root hydraulic properties at each root position (only shown if significant, *p* < 0.05). *p*-values in (**C**) and (**D**) indicate the significance of cultivar effects (S. Dickkopf vs. Tommi) on *k*_x_ and *K*_r_ at each root position, based on a Wilcoxon test.

*k*_x_ and *k*_r_ exhibited opposite patterns along the root axis: averaged across all cultivars, *k*_x_ decreased almost 50% from the root base toward the tip (1.40 to 0.71 m^4^ MPa^-1^ s^-1^ × 10^-9^), whereas *k*_r_ increased around 35% (from 0.74 to 1.0 m MPa^-1^ s^-1^ × 10^-7^). Averaged across root positions, *k*_x_ also declined significantly with release year (1.20 to 0.79 m^4^ MPa^-1^ s^-1^ × 10^-9^ from the oldest to the most modern cultivar), while *k*_r_ showed no significant trend. However, when *k*_r_ was scaled to account for root diameter—and thus surface area—differences along the root and across cultivars (obtaining the radial conductance *K*_r_, Materials & Methods), a significant decreasing trend with release year was detected (*p* < 0.05, Table 1).

To further illustrate the spatial dynamics along the root, we compared *k*_x_ and *K*_r_ between the oldest and most modern cultivars, which were expected to show the strongest contrast according to previous studies (Baca Cabrera *et al*., 2025). For this, we used the distance from the root base where the different root segments were collected (∼0, 10, and 20 cm; Materials & Methods). Along the root, *k*_x_ was systematically lower in the modern cultivar compared to the oldest (∼25–42%), whereas *K*_r_ showed the opposite trend, albeit less pronounced (Fig. 4C–D). This opposite behaviour of *k*_x_ and *K*_r_ was consistent with patterns typically assumed in mechanistic models and observed in previous empirical studies (see Discussion).

## Discussion

### Root position and cultivar release year drive variation in root anatomy

This study analyzed the variation in root anatomy along crown roots of six German wheat cultivars spanning more than 100 years of breeding history. Across all cultivars, anatomical tissue size decreased from the base toward the tip, consistent with spatial gradients reported for both seminal and crown roots under different cultivars and environmental conditions (Bramley *et al*., 2009; Hendel *et al*., 2021; Yang *et al*., 2024; Guhr *et al*., 2025).

Aerenchyma was generally scarce in these crown roots. The proportion of cortex occupied by aerenchyma lacunae declined from basal to distal positions, yet absolute values remained low (<6%) and many samples lacked aerenchyma entirely. Although aerenchyma is a typical anatomical feature in seminal and nodal roots of wheat—particularly under hypoxic conditions—it can also occur under non-stress conditions (Thomson et al., 1990).

These spatial gradients were further reflected in the principal component analysis (PCA), with basal, mid, and distal positions clearly separated along PC1, dominated by tissue size. Cultivar clustering was less pronounced, but the oldest cultivar (S. Dickkopf) consistently separated from the others, characterized by larger root area and thicker cortex. Notably, S. Dickkopf also displayed higher root axis number than the modern cultivars (Baca Cabrera *et al*., 2025), consistent with an overall larger root system. While previous studies reported a breeding-driven reduction of root system size in wheat (Fradgley *et al*., 2020; Maqbool *et al*., 2022), its impact on anatomy remained unclear. The reductions in tissue size observed here suggest that cultivar differences associated with release year may have influenced root anatomical investment, pointing to a linkage between anatomical and morphological adjustments (Guhr *et al*., 2025). Furthermore, loadings in the PCA revealed three coherent anatomical trait groups: (i) cortex traits (thickness and cell diameter), (ii) tissue size traits and (iii) aerenchyma. This grouping is biologically meaningful: variation in tissue areas reflects normal maturation processes along the root axis (Clément *et al*., 2022; Petrova *et al*., 2023), whereas aerenchyma is associated with stress responses in cereals (Yamauchi *et al*., 2013; Lynch *et al*., 2021). By contrast, cortex traits varied less strongly across root positions and cultivars, which could explain their distinct clustering.

Among all traits, stele area and total metaxylem area showed particularly pronounced changes. Variation in metaxylem area was primarily driven by differences in vessel number rather than diameter. A similar metaxylem area decrease along seminal roots (Hendel *et al*., 2021) and reductions in both metaxylem area and number along crown roots (Wu *et al*., 2011; Kadam *et al*., 2015) have been reported previously in wheat. Unlike seminal roots, which typically contain a single central metaxylem, crown roots can produce multiple vessels (Watt *et al*., 2008; Jones *et al*., 2025). This structural flexibility allows adjustment of total metaxylem area primarily through vessel number rather than excessive diameter increases, possibly balancing transport efficiency and cavitation risk (Harrison Day *et al*., 2023), and the observed differences among cultivars could reflect subtle variation in root plasticity (Karlova *et al*., 2021). Interestingly, in all cultivars studied here, a similar spatial pattern in metaxylem number—with a decrease from the root base toward the tip—was observed, which became more pronounced in modern cultivars, as metaxylem number and area were significantly affected by both root position and cultivar release year (Fig 2B, Table 1).

Furthermore, allometric ratio analyses revealed interactions between cultivar release year and root position for XCS and XRR, two established metrics for analyzing anatomical investment in grasses (Yamauchi *et al*., 2021). Both ratios increased significantly with cultivar release year at the distal position but not at the base or in the mid-root, indicating a shift in anatomical composition along the root axis, driven mainly by changes in vascular tissue areas. These allometric shifts were more pronounced in modern than in older cultivars, which could, at least in part, reflect differences among cultivars in maturation rate along the root axis. While distinctions between isometric and allometric scaling in wheat crown roots remain largely unexplored, the phenotyping approach applied here provides a robust framework to investigate how genotypic differences shape these anatomical trade-offs.

Finally, the cultivars showed consistent increases in apoplastic barriers along the root axis. While almost no hypodermal modifications were detected at distal positions, a suberized exodermis became progressively more frequent towards the base. A similar spatial pattern was observed for multilayer cortical sclerenchyma (MCS) and for the combination of MCS and suberized exodermis. Such apoplastic barriers are common in cereal crops (Schneider, 2022; Liu and Kreszies, 2023; Jones *et al*., 2025) and also occurred in the investigated wheat cultivars, which were grown under non-stress field conditions (Baca Cabrera *et al*., 2025; Behrend *et al*., 2025, Preprint). Furthermore, root cortical senescence (RCS) was practically absent, likely reflecting the non-stress conditions under which the plants were grown, but it may also indicate that the sampled crown roots had not yet reached the developmental stage at which RCS typically occurs (Schneider *et al*., 2017). Cultivar effects on hypodermal modifications were minor compared with the strong spatial gradients, and no clear patterns were evident regarding the influence of release year. Overall, these spatial gradients reflect typical cortical maturation, including structural and functional modifications mediated by lignin and suberin deposition and programmed cell death in defined spatial and temporal patterns (Jones *et al*., 2025).

### Implications of anatomical variation for root hydraulics and water uptake capacity

The anatomical patterns observed here are expected to have direct consequences for root hydraulic properties, which we modelled using GRANAR–MECHA. Radial conductivity (*k*_r_) and axial conductance (*k*_x_) displayed opposite spatial trends: *k*_r_ increased from the base to distal positions, whereas *k*_x_ declined. These trends are consistent with experimental data on wheat (Bramley *et al*., 2009) and barley (Knipfer and Fricke, 2011), as well as with common model parameterizations (Doussan *et al*., 1998; Meunier *et al*., 2018). Across cultivars, *k*_x_ decreased by approximately 50% and *K*_r_ (*k*_r_ scaled by root diameter) increased by about 35% from basal to distal positions. These gradients, although much smaller than the order-of-magnitude variation often assumed in models (Doussan *et al*., 1998), provide complementary information, as all sampled sections corresponded to mature root segments with fully developed metaxylem vessels and an endodermis containing a Casparian strip. This highlights variation in more mature portions of the root that is not captured by typical model parameterizations. This is particularly notable, because empirical studies have mostly focused on the pronounced spatial gradients from the root tip (meristematic and elongation zones) toward the base (Bramley *et al*., 2009; Knipfer and Fricke, 2011), whereas our study indicates that spatial gradients also persist within mature tissues—i.e. regions beyond the elongation zone, where primary growth is complete and the stele and cortex are fully differentiated.

Spatial variation in *k*_r_ was primarily driven by the development of apoplastic barriers, particularly suberization of the exodermis. The role of the Casparian strip and suberin lamellae in restricting radial transport is well established (Peterson *et al*., 1993; Steudle *et al*., 1993; Frensch *et al*., 1996; Geldner, 2013). Apoplastic barrier formation has been shown to substantially reduce radial water flow in wheat and rice (Lu and Fricke, 2023; Song *et al*., 2023) and our observations are consistent with these findings. Furthermore, in our dataset aerenchyma formation increased toward the base and may have contributed modestly to reduced *k*_r_. However, overall aerenchyma proportions were low, so its impact remained minor relative to apoplastic barriers. Cortical traits such as cell diameter and number also varied only slightly across cultivars and positions, further suggesting that their influence on *k*_r_ was small compared with apoplastic barrier effects.

The *k*_r_ values estimated here fell well within the range of experimental measurements for wheat root segments (0.53–1.84 m MPa⁻¹ s⁻¹ × 10⁻⁷; Bramley et al., 2009), supporting the robustness of our simulations. In contrast, *k*_x_ values were more than one order of magnitude higher than those reported for wheat (Bramley *et al*., 2007, 2009) and other grasses (Baca Cabrera *et al*., 2024), but they were consistent with typical model parameterization values (Doussan *et al*., 1998). This discrepancy likely resulted from the use of the Hagen–Poiseuille law, which can overestimate *k*_x_ by up to one order of magnitude (Boursiac *et al*., 2022*a*), as it does not account for limitations due to embolism (Harrison Day *et al*., 2023) or for the intrinsic properties of the xylem vessel network—such as vessel length, branching, connectivity, and the overall topology of the root system (Barry *et al*., 2025).

Building on these spatial gradients, we assessed whether cultivar release year influenced root hydraulic properties. No significant effect of release year on *k*_r_ was detected, consistent with the absence of systematic differences in apoplastic barrier formation. However, when scaling *k*_r_ by root diameter to obtain *K*_r_, a significant decrease with release year emerged, supporting the view that diameter is a useful proxy for radial transport capacity (Heymans, 2022). In contrast, variation in *k*_x_ was more strongly associated with cultivar differences, with a clear decline in modern cultivars. Across all cultivars, the spatial decline in *k*_x_ was linked to reductions in metaxylem number and area from the base toward the tip, as reported previously for wheat (Bramley *et al*., 2009). Reduced metaxylem area and vessel number have been linked to reduced water uptake in wheat (Richards and Passioura, 1989; Yang *et al*., 2024) and may reflect a shift toward more conservative water-use strategies. Furthermore, the trends in root hydraulic properties with cultivar release year were more pronounced in *k*_x_ than in *k*_r_, highlighting the importance of axial conductance in shaping water uptake capacity. While *k*_r_ has historically been considered the main limitation to root water transport (Frensch and Steudle, 1989; Bramley *et al*., 2009), recent studies emphasize that *k*_x_ can also critically constrain flow, particularly when metaxylem development or vessel connectivity is limited (Bouda *et al*., 2018; Barry *et al*., 2025). Consistently, Li *et al*. (2024) found that smaller xylem vessels were associated with higher water-use efficiency, while higher cortical thickness enhanced water storage capacity in dryland wheat cultivars—together indicating that both axial and radial traits can modulate root water uptake.

Moreover, the observed decrease in *k*_r_, *K*_r_, and *k*_x_ with cultivar release year along crown roots suggests a corresponding decline in whole-root system conductance (*K*_rs_). This aligns with our previous findings of reduced *K*_rs_ across the same historical wheat cultivars (Baca Cabrera *et al*., 2025). In that earlier study, the decline was primarily attributed to a reduced number of root axes—i.e. fewer “pipes” for water transport—but we also hypothesized that anatomical differences could contribute to that decrease. The decrease in root hydraulic properties with cultivar release year observed here supports this interpretation. To test this further, we simulated *K*_rs_ for the most contrasting cultivars (oldest vs. most modern) with CPlantBox (Giraud *et al*., 2023), using the *k*_r_, and *k*_x_ values determined here and the root architectures as in Baca Cabrera *et al*. (2025). The model reproduced the observed differences in *K*_rs_ between cultivars and closely matched the measured values from the previous study (Figure S4). Notably, these differences emerged even before tillering, indicating that they were driven not only by root axis number but also by segment-scale anatomical properties affecting *k*_r_ and *k*_x_.

The observed cultivar differences in root hydraulic properties should be interpreted considering several methodological aspects. GRANAR–MECHA offers limited flexibility to represent the full diversity of apoplastic barriers, and features such as multiseriate cortical sclerenchyma (MCS) are not explicitly implemented. Since MCS occurred consistently across cultivars, this omission is unlikely to bias comparative trends, although it may affect absolute *k*_r_ values, as MCS influences radial water fluxes, even though its function as a barrier is not fully understood (Schneider, 2022). Additionally, our scoring of the endodermis required a choice between “Casparian strip only” or “fully suberized” depending on tissue maturity—a necessary simplification that affects *k*_r_ estimates but best reflects the observed anatomical state. In addition, cultivar differences could partly arise from variation in cell-scale hydraulic properties, such as membrane permeability or aquaporin (AQP) activity, for which data remain limited. Furthermore, *k*_x_ was quantified solely from metaxylem vessel number and area, excluding protoxylem, which likely contributes minimally once metaxylem elements are functional (Wu *et al*., 2011). Finally, we assumed a constant AQP contribution across cultivars and root sections. Although this choice allowed us to determine anatomical effects on *k*_r_, it does not fully capture known variation in AQP expression, which in wheat exhibits strong diurnal regulation and declines under stress (Lu and Fricke, 2023). Future experimental and modelling efforts should integrate dynamic AQP regulation to better represent root system responses under fluctuating environmental conditions.

Taken together, our results suggest that cultivar-related shifts over release years have been accompanied by coordinated anatomical changes leading to lower axial and radial conductance, likely translating into a reduced whole-root water uptake capacity. Such shifts may reflect a trend toward more conservative water-use behavior in modern cultivars, which can become beneficial under limited water availability. This pattern could, at least partially, be associated with a faster maturation of root tissues in modern cultivars, leading to an earlier development of anatomical barriers and a corresponding reduction in radial conductance. In addition to these physiological insights, the *k*_r_ and *k*_x_ values quantified beyond the maturation zone provide key parameters for modelling efforts that integrate root architecture and hydraulics to predict water uptake across contrasting genotypes and environmental conditions.

### Conclusions and future perspectives

This study revealed clear longitudinal and release year gradients in root anatomical and hydraulic traits in German winter wheat cultivars spanning over 100 years of breeding. Across cultivars, anatomical traits varied strongly along the crown root axis, with progressive differentiation of cortical and vascular tissues, increased apoplastic barrier development, and contrasting trends in *k*_r_, *K*_r_, and *k*_x_. A clear trend with cultivar year of release was also observed, with modern cultivars showing smaller tissue dimensions and fewer metaxylem vessels, together indicating a coordinated shift in anatomical investment. These differences translated into lower radial and axial conductance, consistent with the notion that recent breeding has favored more conservative root water uptake strategies.

Beyond these findings, this study highlights the potential of the presented approach for phenotyping root hydraulic properties. The integration of RAT with the GRANAR–MECHA modelling framework enabled detailed, spatially explicit estimation of segment-scale hydraulic properties from field-grown wheat cultivars, effectively linking anatomical measurements to functional water uptake capacity. Expanding this framework to larger genotype panels and diverse environments could help identify hydraulic traits associated with drought tolerance, resource-use efficiency, and climate resilience. Ultimately, combining anatomical–hydraulic modelling with high-throughput imaging and root architectural simulations could establish a systematic approach for assessing root hydraulics in cereals and translating anatomical and physiological insights into relevant traits for breeding.

## Supplementary data

**Table S1**: Scoring system for hypodermal modifications in crown root cross-sections

**Table S2:** Subcellular hydraulic parameterization used in MECHA

**Table S3:** Root anatomical and hydraulic traits and tissue ratios of winter wheat (*T. aestivum* L.)

**Figure S1:** Meteorological conditions during the field experiment

**Figure S2:** Cross-section images of winter wheat (*T. aestivum* L.) crown roots

**Figure S3:** Example of reconstructed root cross-sections using GRANAR

**Figure S4:** Cultivar effect on the development of root system conductance (*K*_rs_) over plant age

## Acknowledgments

The authors thank Valentin Couvreur and Marco D’Agostino (Earth and Life Institute, UC-Louvain) for helpful discussions on the apoplastic barrier parameterization in the MECHA model.

## Author contributions

JCBC: Conceptualization, Data Curation, Formal Analysis, Investigation, Visualization, Writing - original draft, Writing - review and editing

DHJ: Data Curation, Investigation, Methodology, Writing - review and editing JV: Funding Acquisition, Writing - review and editing

DB: Resources, Writing - review and editing

HS: Funding Acquisition, Resources, Writing - review and editing

GL: Formal Analysis, Funding Acquisition, Writing - review and editing

## Data availability

Data supporting the findings of this study are available within the paper, within its Supplementary data (Tables S1–S3, Figures S1–S4) or on request.

