## Supplementary Tables S1-S3 for "Root anatomical gradients and cultivar differences underlie variation in root hydraulic properties in German winter wheat"

**Table S1**: Scoring system for hypodermal modifications in crown root cross-sections. A score of 1–5 was assigned according to described optical features. Example images for each score are shown for reference. All imaging was performed at the Leibniz Institute of Plant Genetics and Crop Plant Research (IPK), Gatersleben.

| | | |
|---|---|---|
| **Type 1**<br><br>No<br>modif cations | Images were assigned type 1 hypodermis if no clear dif erences were observed between inner and outer cortical cell f le layers in cell wall brightness, brightness pattern, cell size, or density. | 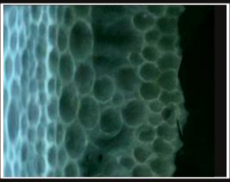 |
| **Type 2**<br><br>Polar<br>exodermis | Images were assigned type 2 hypodermis where there is a visible dif erence in cell wall f uorescence pattern in the exodermal layer compared to inner layers, but cell size and layer density consistent with mesodermis | 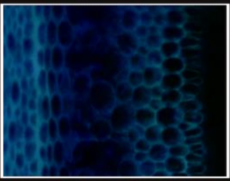 |
| **Type 3**<br><br>MCS like<br>hypodermis | Images were assigned type 3 hypodermis where there are multiple layers of brighter (lignif ed) cell walls in the hypodermis, with distinctly smaller cells than in the mesodermis | 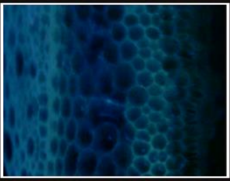 |
| **Type 4**<br><br>Type 2 and<br>3 present | Images were assigned type 4 hypodermis where the structures described in type 2 (polar exodermis) and 3 (multiple layers of small and bright cell walls) were observed together | 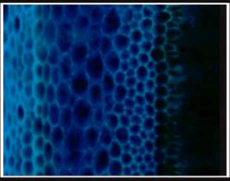 |
| **Type 5**<br><br>Cortical<br>senescence | Images were assigned type 5 hypodermis when the majority of cortical cell f le layers were senesced, leading to total or near total loss of visibility in distinct cell walls. Endodermis was still present. | 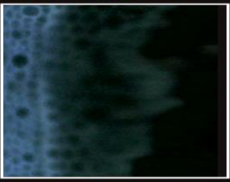 |

**Table S2**: Subcellular hydraulic parameterization used in MECHA. Default hydraulic parameters were used for all simulations, as described in Couvreur *et al*. (2018, Table 2) and Heymans *et al*. (2019, Materials and Methods).

| **Parameter** | **Value** | **Unit** |
|---|---|---|
| Cell wall hydraulic conductivity | $2.8 \cdot 10^{-9}$ | $m^2\ MPa^{-1}\ s^{-1}$ |
| Hydraulic conductivity of suberized and lignified cell walls | Hydrophobic | - |
| Plasma membrane intrinsic hydraulic conductivity ($k_m$) | $3.5 \cdot 10^{-8}$ | $m\ MPa^{-1}\ s^{-1}$ |
| Contribution of aquaporins to plasma membrane hydraulic conductivity ($k_{AQP}$) | $5.0 \cdot 10^{-7}$ | $m\ MPa^{-1}\ s^{-1}$ |
| Conductance of plasmodesmata per unit membrane surface ($k_{PD}$) | $2.4 \cdot 10^{-7}$ | $m\ MPa^{-1}\ s^{-1}$ |
| Protoplast permeability ($L_{pc}$) | $7.74 \cdot 10^{-7}$ | $m\ MPa^{-1}\ s^{-1}$ |

**Table S3**: Root anatomical and hydraulic traits and tissue ratios of winter wheat (*T. aestivum* L.). Data correspond to mean ± SE ($n$ = 32) for six German winter wheat cultivars sampled at three different root positions.

| parameter | root position | S. Dickkopf (1895) | SG v. Stocken (1920) | Heines II (1940) | Jubilar (1961) | Okapi (1978) | Tommi (2002) |
|---|---|---|---|---|---|---|---|
| **root area ($mm^2$)** | base | 0.53 ± 0.03 | 0.47 ± 0.02 | 0.49 ± 0.03 | 0.48 ± 0.02 | 0.51 ± 0.02 | 0.49 ± 0.02 |
| | mid-root | 0.47 ± 0.02 | 0.38 ± 0.02 | 0.39 ± 0.03 | 0.37 ± 0.01 | 0.4 ± 0.02 | 0.38 ± 0.02 |
| | near-tip | 0.45 ± 0.02 | 0.35 ± 0.02 | 0.35 ± 0.02 | 0.32 ± 0.02 | 0.37 ± 0.02 | 0.35 ± 0.02 |
| **metaxylem area ($mm^2$)** | base | 0.0016 ± 7e-05 | 0.0015 ± 6e-05 | 0.0017 ± 9e-05 | 0.0016 ± 9e-05 | 0.0016 ± 8e-05 | 0.0015 ± 8e-05 |
| | mid-root | 0.0015 ± 7e-05 | 0.0013 ± 7e-05 | 0.0013 ± 7e-05 | 0.0015 ± 9e-05 | 0.0015 ± 9e-05 | 0.0012 ± 7e-05 |
| | near-tip | 0.0015 ± 1e-04 | 0.0011 ± 6e-05 | 0.0013 ± 6e-05 | 0.0015 ± 8e-05 | 0.0015 ± 9e-05 | 0.0012 ± 1e-04 |
| **metaxylem number** | base | 5.5 ± 0.2 | 5.1 ± 0.2 | 5 ± 0.2 | 5.2 ± 0.2 | 5.2 ± 0.2 | 4.9 ± 0.2 |
| | mid-root | 4.9 ± 0.2 | 4.7 ± 0.1 | 4.8 ± 0.2 | 4.6 ± 0.1 | 4.4 ± 0.2 | 4.5 ± 0.2 |
| | near-tip | 4.7 ± 0.2 | 4.2 ± 0.2 | 4.2 ± 0.2 | 4 ± 0.2 | 3.6 ± 0.2 | 3.9 ± 0.2 |
| **total metaxylem area ($mm^2$)** | base | 0.0089 ± 6e-04 | 0.0074 ± 3e-04 | 0.0083 ± 5e-04 | 0.0081 ± 5e-04 | 0.008 ± 4e-04 | 0.0071 ± 4e-04 |
| | mid-root | 0.0072 ± 4e-04 | 0.0058 ± 3e-04 | 0.0065 ± 5e-04 | 0.0068 ± 4e-04 | 0.0066 ± 4e-04 | 0.0051 ± 3e-04 |
| | near-tip | 0.0064 ± 4e-04 | 0.0047 ± 3e-04 | 0.0053 ± 3e-04 | 0.0057 ± 4e-04 | 0.0053 ± 4e-04 | 0.0044 ± 3e-04 |
| **cortical cell diameter (mm)** | base | 0.028 ± 9e-04 | 0.027 ± 6e-04 | 0.026 ± 6e-04 | 0.027 ± 4e-04 | 0.027 ± 4e-04 | 0.027 ± 6e-04 |
| | mid-root | 0.03 ± 0.001 | 0.027 ± 7e-04 | 0.028 ± 9e-04 | 0.027 ± 6e-04 | 0.028 ± 7e-04 | 0.027 ± 6e-04 |
| | near-tip | 0.031 ± 0.001 | 0.029 ± 0.001 | 0.028 ± 7e-04 | 0.028 ± 8e-04 | 0.03 ± 8e-04 | 0.028 ± 8e-04 |

| parameter | root position | S. Dickkopf (1895) | SG v. Stocken (1920) | Heines II (1940) | Jubilar (1961) | Okapi (1978) | Tommi (2002) |
|---|---|---|---|---|---|---|---|
| **cortical cell number** | base | 8.6 ± 0.2 | 8.7 ± 0.2 | 8.9 ± 0.2 | 8.6 ± 0.1 | 9.2 ± 0.2 | 8.5 ± 0.2 |
| | mid-root | 7.9 ± 0.1 | 8 ± 0.2 | 7.8 ± 0.2 | 7.9 ± 0.1 | 7.9 ± 0.1 | 7.8 ± 0.1 |
| | near-tip | 7.7 ± 0.3 | 7.2 ± 0.2 | 7.4 ± 0.2 | 7.4 ± 0.2 | 7.2 ± 0.1 | 7.3 ± 0.2 |
| **total cortex thickness (mm)** | base | 0.24 ± 0.007 | 0.23 ± 0.005 | 0.23 ± 0.007 | 0.23 ± 0.004 | 0.24 ± 0.006 | 0.23 ± 0.006 |
| | mid-root | 0.23 ± 0.008 | 0.22 ± 0.006 | 0.22 ± 0.008 | 0.21 ± 0.004 | 0.22 ± 0.005 | 0.21 ± 0.004 |
| | near-tip | 0.23 ± 0.008 | 0.21 ± 0.008 | 0.21 ± 0.006 | 0.2 ± 0.007 | 0.21 ± 0.005 | 0.21 ± 0.007 |
| **stele area ($mm^2$)** | base | 0.071 ± 0.004 | 0.06 ± 0.003 | 0.069 ± 0.004 | 0.069 ± 0.003 | 0.063 ± 0.003 | 0.066 ± 0.003 |
| | mid-root | 0.055 ± 0.002 | 0.046 ± 0.002 | 0.049 ± 0.003 | 0.052 ± 0.002 | 0.046 ± 0.002 | 0.046 ± 0.002 |
| | near-tip | 0.051 ± 0.003 | 0.037 ± 0.001 | 0.039 ± 0.002 | 0.041 ± 0.002 | 0.039 ± 0.002 | 0.036 ± 0.002 |
| **aerenchyma number** | base | 3.8 ± 0.7 | 5.7 ± 0.7 | 5.3 ± 0.9 | 8.8 ± 1 | 7.9 ± 1 | 4.9 ± 0.9 |
| | mid-root | 2.1 ± 0.5 | 3.6 ± 0.7 | 5.6 ± 1 | 6.7 ± 2 | 5.9 ± 1 | 3.3 ± 0.9 |
| | near-tip | 2 ± 0.6 | 1.8 ± 0.5 | 2.1 ± 0.6 | 3.3 ± 0.9 | 2.2 ± 0.6 | 1.7 ± 0.5 |
| **aerenchyma proportion** | base | 0.032 ± 0.007 | 0.044 ± 0.006 | 0.042 ± 0.007 | 0.051 ± 0.008 | 0.046 ± 0.007 | 0.034 ± 0.006 |
| | mid-root | 0.025 ± 0.006 | 0.026 ± 0.005 | 0.048 ± 0.009 | 0.041 ± 0.009 | 0.047 ± 0.007 | 0.026 ± 0.008 |
| | near-tip | 0.019 ± 0.006 | 0.015 ± 0.004 | 0.021 ± 0.006 | 0.019 ± 0.005 | 0.018 ± 0.005 | 0.012 ± 0.003 |
| **$k_x$ ($m^4$ $MPa^{-1}$ $s^{-1}$ × $10^{-9}$)** | base | 1.6 ± 0.2 | 1.2 ± 0.08 | 1.5 ± 0.2 | 1.4 ± 0.2 | 1.3 ± 0.1 | 1.2 ± 0.1 |
| | mid-root | 1.1 ± 0.1 | 0.75 ± 0.08 | 0.97 ± 0.1 | 1.1 ± 0.1 | 0.98 ± 0.1 | 0.63 ± 0.06 |
| | near-tip | 0.91 ± 0.1 | 0.56 ± 0.06 | 0.64 ± 0.07 | 0.85 ± 0.1 | 0.78 ± 0.09 | 0.57 ± 0.07 |

| parameter | root position | S. Dickkopf (1895) | SG v. Stocken (1920) | Heines II (1940) | Jubilar (1961) | Okapi (1978) | Tommi (2002) |
|---|---|---|---|---|---|---|---|
| | base | 0.8 ± 0.07 | 0.77 ± 0.05 | 0.71 ± 0.07 | 0.71 ± 0.07 | 0.67 ± 0.06 | 0.78 ± 0.07 |
| **$k_r$ (m MPa$^{-1}$ s$^{-1}$ × 10$^{-7}$)** | mid-root | 0.98 ± 0.05 | 0.94 ± 0.05 | 0.85 ± 0.08 | 0.94 ± 0.09 | 0.95 ± 0.07 | 0.91 ± 0.06 |
| | near-tip | 1.1 ± 0.1 | 1.1 ± 0.05 | 1 ± 0.07 | 1.1 ± 0.05 | 1 ± 0.05 | 1 ± 0.05 |
| | base | 2 ± 0.2 | 1.8 ± 0.1 | 1.7 ± 0.2 | 1.7 ± 0.2 | 1.6 ± 0.1 | 1.9 ± 0.2 |
| **$K_r$ (m$^3$ MPa$^{-1}$ s$^{-1}$ × 10$^{-7}$)** | mid-root | 2.3 ± 0.1 | 2 ± 0.1 | 1.8 ± 0.2 | 2 ± 0.2 | 2.1 ± 0.1 | 1.9 ± 0.1 |
| | near-tip | 2.5 ± 0.2 | 2.1 ± 0.09 | 2 ± 0.1 | 2.2 ± 0.1 | 2.2 ± 0.1 | 2.1 ± 0.09 |
| | base | 0.024 ± 0.001 | 0.026 ± 0.002 | 0.026 ± 0.002 | 0.023 ± 0.001 | 0.026 ± 0.001 | 0.023 ± 0.001 |
| **XSR** | mid-root | 0.027 ± 0.001 | 0.028 ± 0.002 | 0.028 ± 0.001 | 0.03 ± 0.001 | 0.034 ± 0.002 | 0.027 ± 0.002 |
| | near-tip | 0.03 ± 0.003 | 0.031 ± 0.002 | 0.034 ± 0.002 | 0.037 ± 0.003 | 0.042 ± 0.002 | 0.034 ± 0.002 |
| | base | 7 ± 0.4 | 7 ± 0.3 | 6.2 ± 0.3 | 6 ± 0.2 | 7.2 ± 0.2 | 6.6 ± 0.3 |
| **CSR** | mid-root | 7.7 ± 0.3 | 7.3 ± 0.3 | 7.2 ± 0.3 | 6.3 ± 0.2 | 7.9 ± 0.3 | 7.3 ± 0.3 |
| | near-tip | 8.3 ± 0.4 | 8.3 ± 0.5 | 8.3 ± 0.4 | 7.2 ± 0.4 | 9 ± 0.4 | 8.9 ± 0.4 |
| | base | 0.0031 ± 2e-04 | 0.0033 ± 2e-04 | 0.0038 ± 3e-04 | 0.0034 ± 2e-04 | 0.0032 ± 2e-04 | 0.0031 ± 2e-04 |
| **XRR** | mid-root | 0.0033 ± 2e-04 | 0.0033 ± 2e-04 | 0.0037 ± 3e-04 | 0.0042 ± 3e-04 | 0.0039 ± 2e-04 | 0.0032 ± 2e-04 |
| | near-tip | 0.0035 ± 3e-04 | 0.0036 ± 3e-04 | 0.0038 ± 3e-04 | 0.0048 ± 3e-04 | 0.0043 ± 3e-04 | 0.0037 ± 4e-04 |
| | base | 0.13 ± 0.006 | 0.13 ± 0.004 | 0.14 ± 0.005 | 0.15 ± 0.004 | 0.13 ± 0.003 | 0.14 ± 0.005 |
| **SRR** | mid-root | 0.12 ± 0.004 | 0.12 ± 0.004 | 0.13 ± 0.009 | 0.14 ± 0.005 | 0.12 ± 0.004 | 0.12 ± 0.004 |
| | near-tip | 0.11 ± 0.006 | 0.12 ± 0.007 | 0.11 ± 0.006 | 0.13 ± 0.007 | 0.11 ± 0.004 | 0.11 ± 0.004 |

| parameter | root position | S. Dickkopf (1895) | SG v. Stocken (1920) | Heines II (1940) | Jubilar (1961) | Okapi (1978) | Tommi (2002) |
|---|---|---|---|---|---|---|---|
| **XCS** | base | 0.17 ± 0.02 | 0.19 ± 0.02 | 0.16 ± 0.01 | 0.14 ± 0.007 | 0.18 ± 0.01 | 0.16 ± 0.01 |
| | mid-root | 0.21 ± 0.009 | 0.21 ± 0.02 | 0.21 ± 0.01 | 0.19 ± 0.01 | 0.27 ± 0.02 | 0.2 ± 0.02 |
| | near-tip | 0.25 ± 0.03 | 0.26 ± 0.02 | 0.29 ± 0.02 | 0.28 ± 0.03 | 0.39 ± 0.04 | 0.31 ± 0.03 |
