## Supplementary Figures S1-S4 for "Root anatomical gradients and cultivar differences underlie variation in root hydraulic properties in German winter wheat"

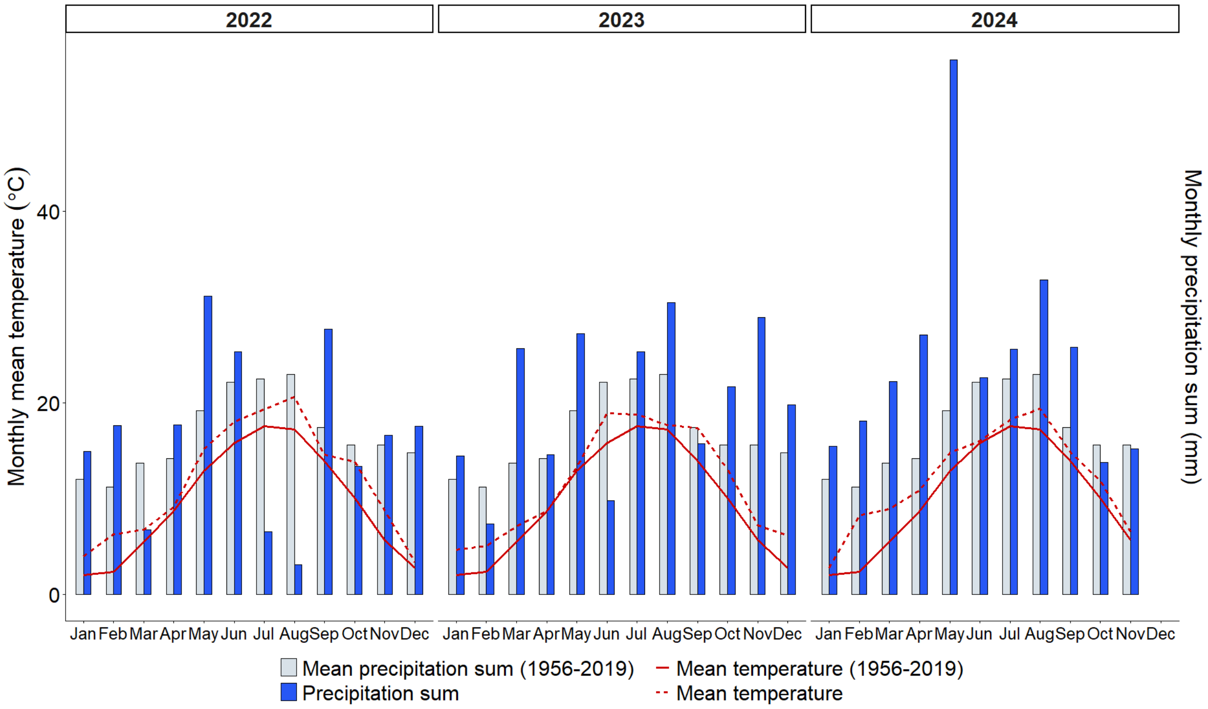


**Figure S1**: Meteorological conditions during the field experiment. Monthly precipitation sums (blue bars) and mean temperatures (dashed red line) during the experimental years (2022, 2023, and 2024), compared with long-term mean precipitation sums (grey bars) and mean temperatures (solid red line) for 1956–2019 at the experimental station Campus Klein-Altendorf.

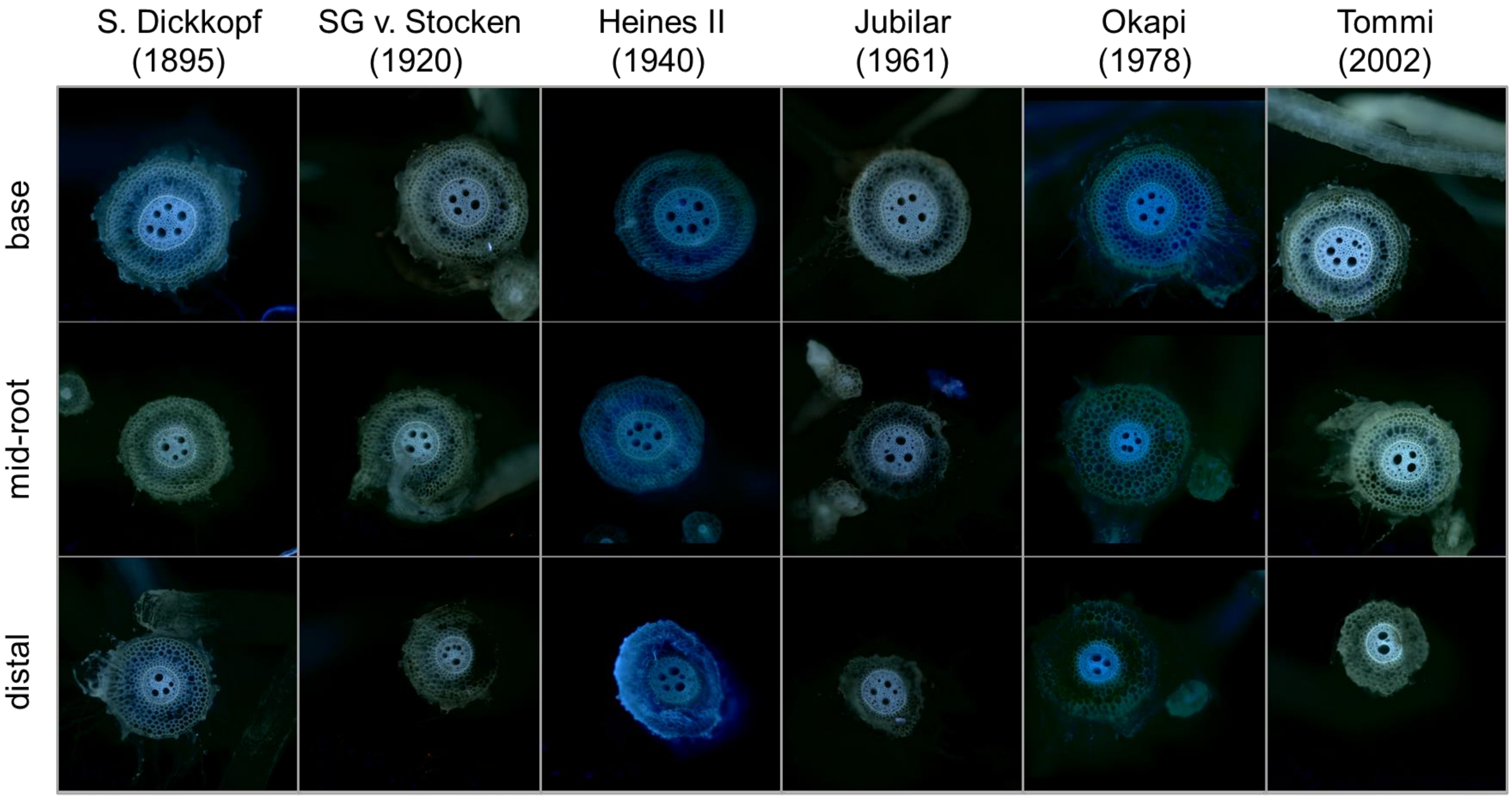


**Figure S2**: Cross-section images of winter wheat (*T. aestivum* L.) crown roots. Example high-quality images obtained with the Rapid Anatomics Tool (Jones *et al*., 2025, Preprint) for six cultivars at three sampling positions (basal, mid-root, distal).

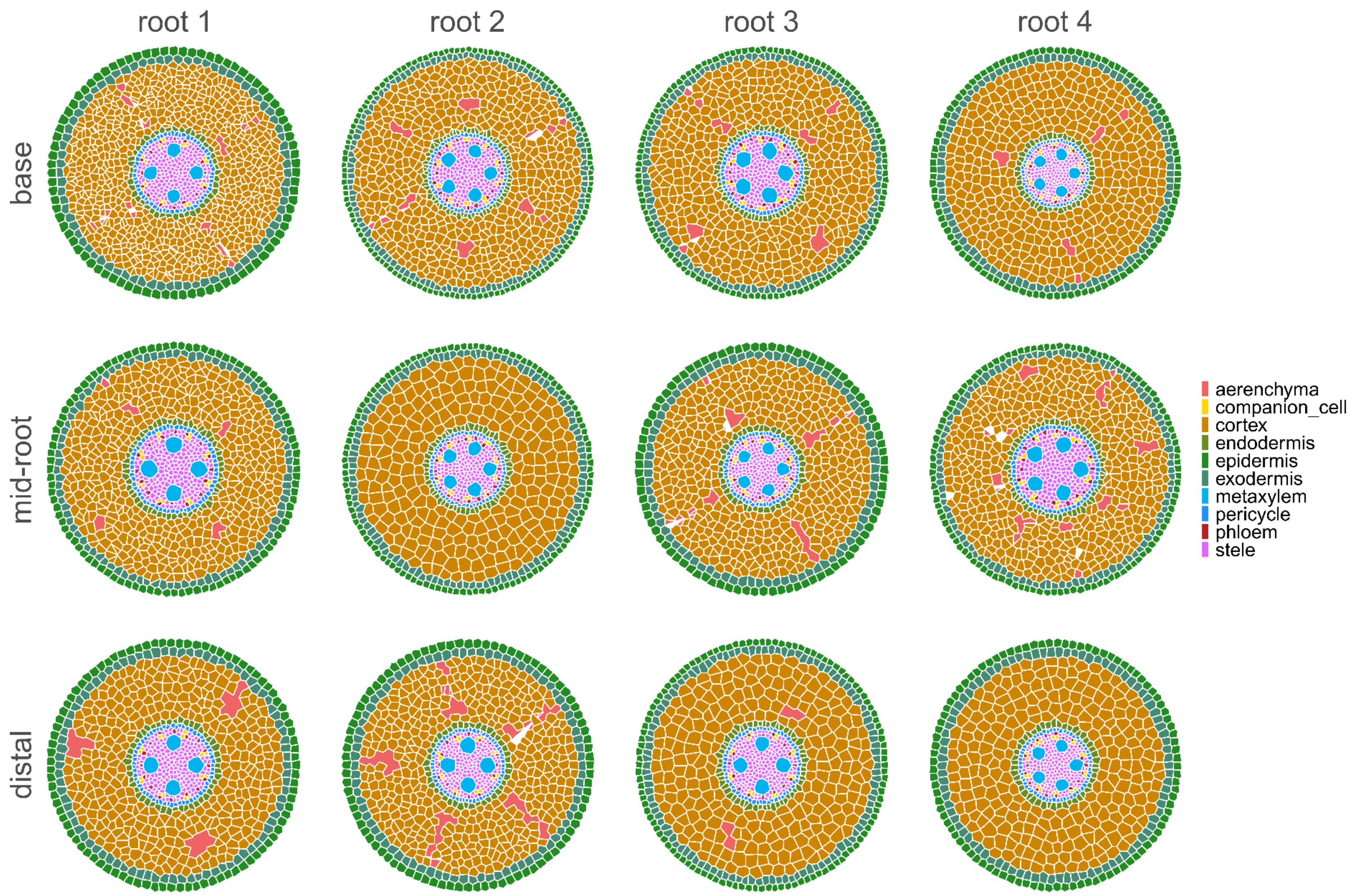


**Figure S3**: Example of reconstructed root cross-sections using GRANAR. The images show samples from the base, mid-root, and distal positions of four different plants sampled in one plot for the German winter wheat cultivar SG v. Stocken.

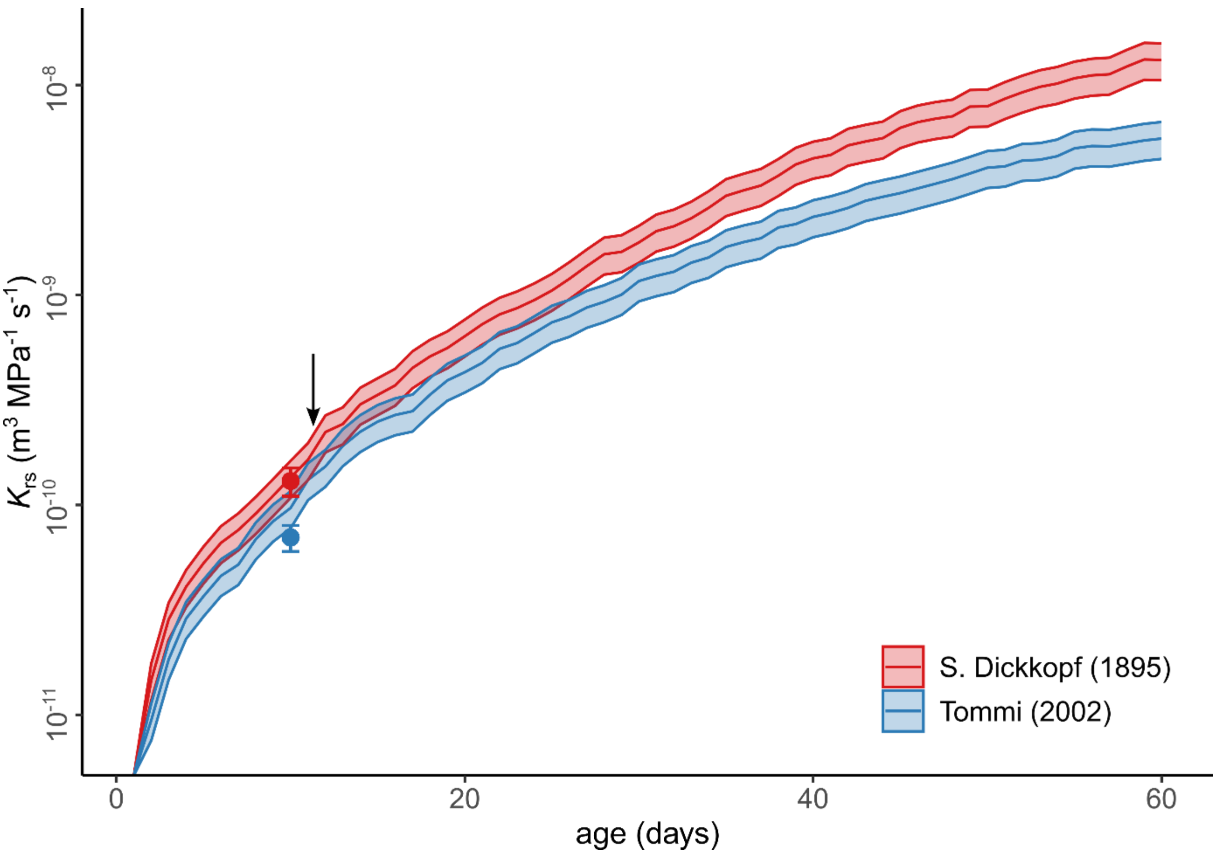


**Figure S4:** Cultivar effect on the development of root system conductance ($K_{rs}$) over plant age. Data are shown for an old (S. Dickkopf, release year 1895) and a modern (Tommi, release year 2002) German winter wheat cultivar (*T. aestivum* L.). Points represent mean ± SE of measured $K_{rs}$ values for the two cultivars (Baca Cabrera *et al.*, 2025). The solid line and shaded area indicate the mean ± SE of simulated $K_{rs}$ (n = 6 simulation runs) using the whole-plant model CPlantBox (Giraud *et al.*, 2023). The model was parameterized using $k_r$ and $k_x$ values estimated at different root positions in this study, combined with root architectures from Baca Cabrera *et al.* (2025). The arrow indicates the onset of tillering, as defined in the model parameterization.
